# Negative interactions and virulence differences drive the dynamics in multispecies bacterial infections

**DOI:** 10.1101/2022.08.10.503476

**Authors:** Désirée A. Schmitz, Richard C. Allen, Rolf Kümmerli

## Abstract

Bacterial infections are often polymicrobial, leading to intricate pathogen-pathogen and pathogen-host interactions. There is increasing interest in studying the molecular basis of pathogen interactions and how such mechanisms impact host morbidity. However, much less is known about the ecological dynamics between pathogens and how they affect virulence and host survival. Here we address these open issues by co-infecting larvae of the insect model host *Galleria mellonella* with one, two, three or four bacterial species, all of which are opportunistic human pathogens. We found that host mortality was always determined by the most virulent species regardless of the number of species and pathogen combinations injected. In certain combinations, the more virulent pathogen simply outgrew the less virulent pathogen. In other combinations, we found evidence for negative interactions between pathogens inside the host, whereby the more virulent pathogen typically won a competition. Taken together, our findings reveal positive associations between a pathogen’s growth inside the host, its competitiveness towards other pathogens, and its virulence. Beyond being generalizable across species combinations, our findings predict that treatments against polymicrobial infections should target the most virulent species to reduce host morbidity, a prediction we validated experimentally.

**Importance:** There is increasing evidence that polymicrobial infections are common and that host morbidity is impacted by interactions between the co-infecting pathogens. In this study, we infected the larvae of the insect host *Galleria mellonella* with four opportunistic human pathogens as mono or multispecies bacterial infections in all possible combinations and followed host survival and pathogen interactions over time. We discovered that species followed the exact same rank order in terms of their virulence, their growth inside the host, and their competitive ability against co-infecting pathogens. As consequence of this consistent rank order, we found that the most virulent species determines host survival dynamics in mixed infections. Beyond revealing a generalizable predictive pattern for virulence outcomes, our findings also predict that treatments should target the most virulent species in polymicrobial infections, a prediction we validated experimentally.

## Introduction

Research over the past decades has revealed that many bacterial infections are polymicrobial (1–4). This can lead to complicated pathogen-pathogen and pathogen-host interactions. The ecological interactions between pathogens can range from competition through commensalism to cooperation, and such interactions can have consequences for the host (5). For example, mutually beneficial cross-feeding between a human commensal bacterium and a pathogen was shown to increase the virulence of the latter, thus exacerbating host morbidity (6). Competition can also enhance virulence as revealed for the pathogen *Pseudomonas aeruginosa* that increases the production of the toxin pyocyanin thereby harming both its competitor *Staphylococcus aureus* and host tissue (7). Interaction patterns can become even more complex in chronic infections, where (co-)evolution can occur between pathogens and between pathogens and the host (8–13). There is increasing awareness that such eco-evolutionary dynamics affect host health and are important to consider in the context of treatment options (5).

For this reason, research on interactions between pathogenic bacteria has flourished in the past years. For example, there is a wealth of work on interactions between *P. aeruginosa and S. aureus*, two of the most troublesome nosocomial pathogens (14–16). This research, often carried out *in vitro*, has successfully identified molecular mechanisms of pathogen interactions and evolutionary patterns of how pathogens adapt to one another. However, we know much less about pathogen-pathogen interactions within hosts and how interactions drive virulence. Moreover, it is often unclear whether insights from a particular pathogen pair are generalizable and hold for other pathogen and strain combinations (17).

Here, we aim to tackle these questions by studying bacterial interactions between four different opportunistic human pathogens in an insect host, the larvae of the greater wax moth *Galleria mellonella*. This model host is suitable for our purpose because it allows for relatively high-throughput experiments where many different pathogen combinations can be tested, and well-defined doses of pathogens can be injected into a larva. Moreover, *G. mellonella* has an innate immune system that resembles the vertebrate innate immune response and lives approximately at human body temperature, which makes it an excellent model organism to study human bacterial pathogenesis (18–21). Regarding the pathogens, our aim was to choose four gram-negative bacterial species that exhibit different levels of virulence in order to test whether generalizable patterns of host survival and pathogen interactions arise across a spectrum of pathogens. We picked the following four opportunistic human pathogens. *Pseudomonas aeruginosa* (P) is a species of high clinical relevance that causes both community- and hospital-acquired infections including skin and wound infections, urinary tract infections, bloodstream infections, and pneumonias (22, 23). *Burkholderia cenocepacia* (B) causes chronic lung infections in immunocompromised patients, e.g. suffering from cystic fibrosis (24). *Klebsiella michiganensis* (K) belongs to the *K. oxytoca* complex (25), which are human pathogens that lead to health care-associated infections as well as to a variety of infections such as diarrhea, bacteremia, and meningitis (26–28). *Cronobacter sakazakii* (C) can also cause bacteremia and meningitis as well as necrotizing enterocolitis and brain abscess/lesions (29). While some of these pathogens can be found together – P and B in lung infections of cystic fibrosis patients (4, 30, 31) and P and K in burn wound infections (32, 33) – co-occurrence for other combinations is rarer. Thus, our model reflects a scenario where different opportunistic pathogens co-incidentally infect the same host without any prior interaction history or coevolution between the co-infecting pathogens. Such a scenario reflects the starting point of many acute and chronic polymicrobial infections.

In a first set of experiments, we aimed to understand the demographics of pathogen-host interactions in single species (mono) infections. For this purpose, we manipulated the injection dose and host age to understand how these factors affect each pathogen’s virulence and mortality for the host. Second, we conducted mixed species infection experiments (double, triple, and quadruple) to assess how polymicrobial infections affect virulence patterns and host survival. Third, we quantified pathogen growth within the host at two time points for all mono and pairwise infections to test whether pathogen load links to virulence. Fourth, we followed changes in pathogen frequencies in mixed infections to derive pathogen interaction patterns. While this approach does not reveal specific molecular mechanisms of pathogen interactions, it allows us to distinguish between negative, neutral, and positive interactions between co-infecting pathogens and how such interactions link to virulence. One key insight from our experiments was that host mortality was always driven by the most virulent species, with the virulence of a pathogen correlating with its growth inside the host and its competitiveness towards the co-infecting species. Based on these insights, we predicted that the most virulent species should be targeted in an infection to reduce host morbidity and mortality. In a follow-up experiment, we validated this prediction by specifically targeting the more virulent *P. aeruginosa* co-infecting the host together with the less virulent *B. cenocepacia*.

## Results

### Injection dose, host age, and species identity determine virulence

To assess how the four different pathogens affect host survival, we exposed *G. mellonella* larvae (last instar larvae, relative age: 5, 10, or 15 days after arrival in our laboratory) to a range of pathogen injection doses (100 to 1 million CFU) for each of the four pathogens (B, C, K, P) and tracked their survival over 48 hours (Fig. 1A and 1B and Fig. S1).

**Figure 1.**
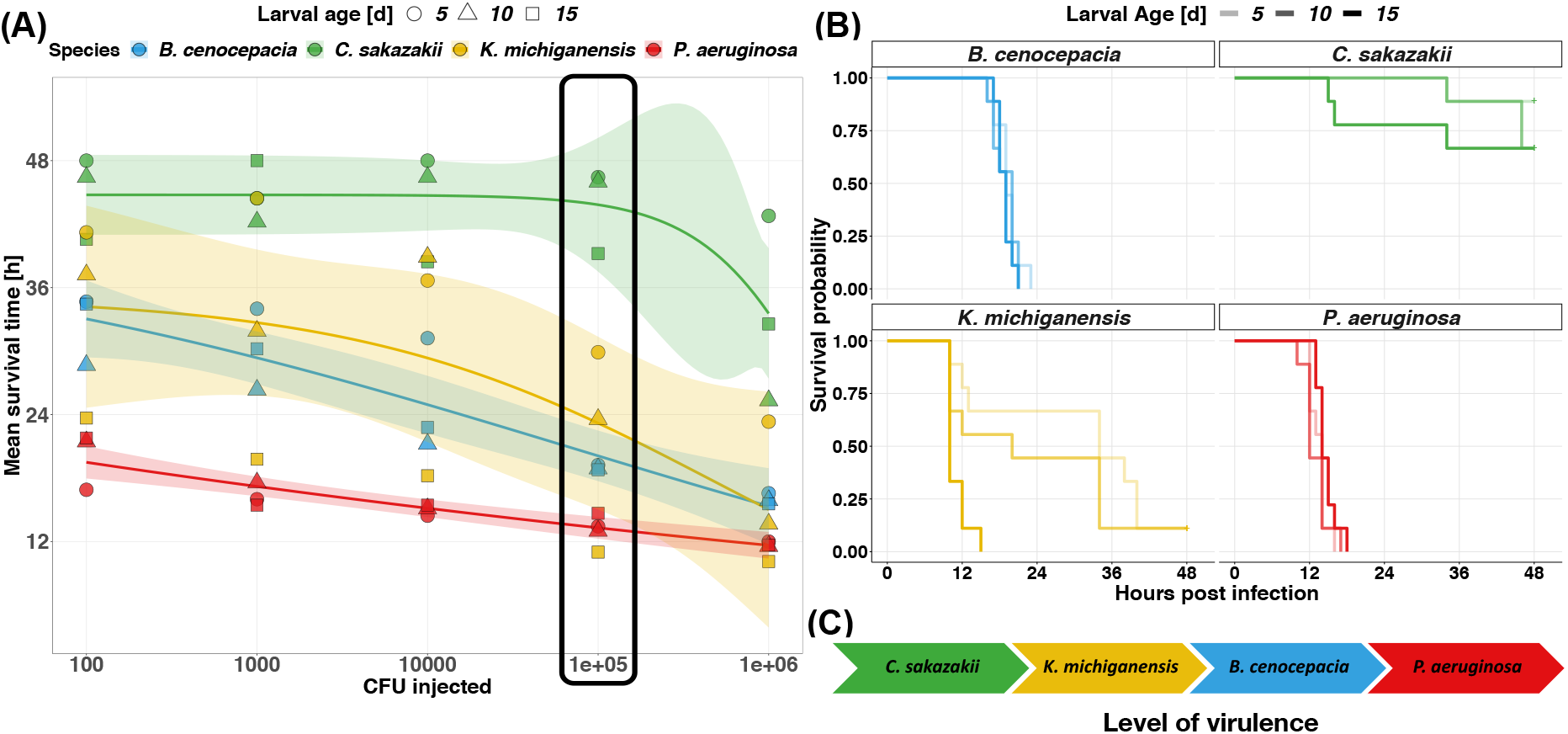
Survival rates of greater wax moth larvae (*G. mellonella*) are affected by the pathogen species they are infected with, injection dose, and larval age. (A) Mean survival time of *G. mellonella* larvae (measured as the area under the full survival curve) as a function of larval age (5, 10, 15 days), injection dose (CFU), and pathogen species (blue: *B. cenocepacia*, green: *C. sakazakii*, yellow: *K. michiganensis*, red: *P. aeruginosa*). The lines and the shaded area (95% CI) depict the relationship between mean survival time (across the different larval ages and replicates) and infection dose for each species. The black box highlights the injection dose chosen for all subsequent experiments. (B) Kaplan Meier survival curves of *G. mellonella* larvae for an injection dose of 10^5^ CFU for each of the four pathogen species. Relative larval age – 5, 10, and 15 days after the arrival in our laboratory – is indicated by increasing line opaqueness. (C) The arrow chart shows the rank order of virulence among the four bacterial pathogens, from lowest to highest based on our experimental data. Data are from three independent experiments, each featuring 10-12 larvae per treatment, resulting in a total of 30-36 larvae per treatment.

We found that the hazard to die differed between the four species (Cox proportional hazard: 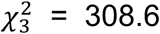, p < 0.0001) and was influenced by significant interactions between species and larval age (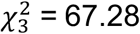, p < 0.0001), and species and injection dose (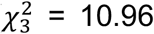, p = 0.0119). When examining these interactions more closely, we observed that the hazard to die grew significantly with increasing larval age for C (z = 2.69, p = 0.0071) and K (z = 6.43, p < 0.0001), but only marginally for B (z = 1.90, p = 0.0573), and not at all for P (z = −0.72, p = 0.471). For the injection dose, we found that higher CFU significantly increased the hazard to die in all four species, but the hazard increase was more pronounced for B and P than for K and C (see Table S1 for the full statistical analysis). The reason for the species-specific relationship between virulence and infection dose stems from the fact that the survival decreased in a log-linear fashion for B and P with higher pathogen doses, whereas C and K only became virulent above a certain threshold injection dose of about 10^4^ CFU for K and 10^5^ CFU for C.

Based on the insights from this experiment, we decided to use an injection dose of 10^5^ CFU for all subsequent experiments. At this injection dose, all four pathogens are virulent with the following order of virulence: *P. aeruginosa* (P) > *B. cenocepacia* (B) > *K. michiganensis* (K) > *C. sakazakii* (C), and with significant host age effects for C and K (Fig. 1C).

### Host mortality in multispecies infections is determined by the most virulent species

Next, we compared host survival between mono and mixed infections. For this, we injected pairwise combinations of our four bacterial species into larvae of *G. mellonella* and observed their survival over 48 h (Fig. 2A). The injection dose was always 10^5^ CFU in total, with equal amounts of each co-injected species. An intuitive expectation is that the virulence in co-infections should be intermediate between the mono infections of the two species. In contrast to this expectation, we found that any mixed infection followed the dynamics of its most virulent species (Fig. 2B). A statistical examination confirmed that larval survival was not different between the mixed infection and the mono infection of the more virulent species in three out of six cases (log-rank test B+K vs. B: p = 0.2459; C+K vs. K: p = 0.9770; C+P vs. P: p = 0.0548). In the remaining three cases, survival in the mixed infection was significantly lower than the mono infection of the more virulent species (B+C vs. B: p = 0.0033; B+P vs. P: p = 0.0230; K+P vs. P: p = 0.0041), but the actual biological differences observed are extremely small (Fig. 2B).

**Figure 2.**
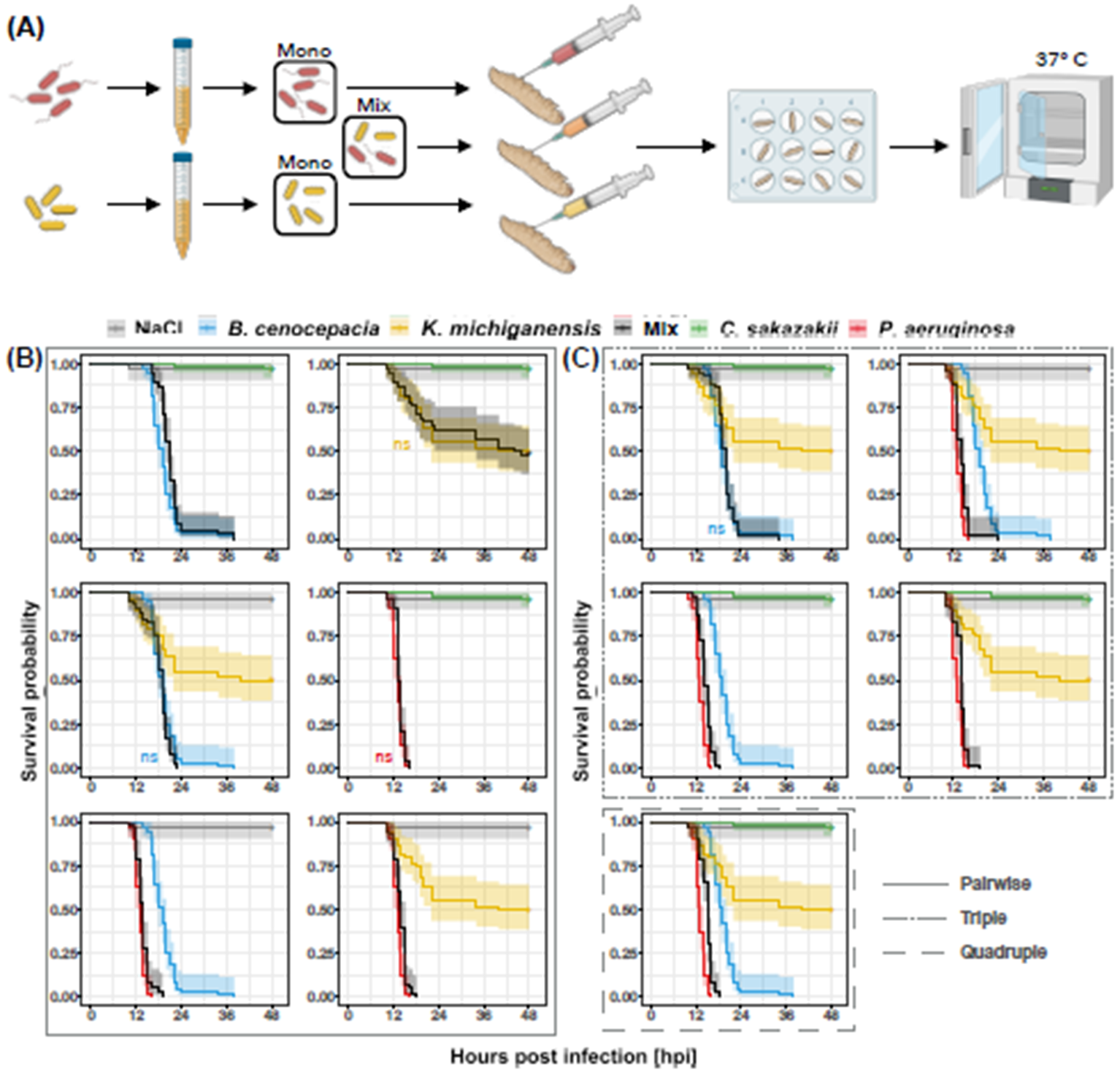
Host survival dynamics of *G. mellonella* larvae of pairwise, triple, and quadruple bacterial infections follow the pattern of the mono infection (colored curves) of the most virulent pathogen in a mix (black curves depict the mix of all pathogens within the respective panel). (A) Schematic overview of the experimental setup with the example of the mono and pairwise infections of *P. aeruginosa* (red) and *K. michiganensis* (yellow). A total of 10^5^ CFU was injected into *G. mellonella* larvae with equal amounts of each species in a mix. Kaplan Meier survival curves of mono versus pairwise (B), triple, and quadruple (C) infections with the shaded area indicating the 95% CI. The denotation ns (non-significance) marks cases for which survival does not significantly differ between one of the mono infections compared to the mixed infection in the same panel. Data are from five independent experiments, each with 10-12 larvae per treatment, resulting in a total of 50-60 larvae per treatment.

We then tested whether the same patterns arise in the four triple and the quadruple infections. As above, we kept the total number of CFU constant at 10^5^ CFU and mixed equal amounts of each pathogen. As for the paired infections, we found that the survival trajectory of larvae infected with three and four pathogens followed the trajectories of the mono infection of the most virulent species in the mix (Fig. 2C). For example, in the triple infection with B, C, and K, larval survival followed the one of the B mono-infection, which is the most virulent of the three species. In the remaining four mixes, the most virulent species was P and in all these cases larval survival followed the one of P mono infections. Statistical analyses revealed subtle but significant differences: mixed infections with P were always slightly less virulent than the mono infections with P (see Table S2 for the full statistical analysis). We can explain this pattern by a density effect: fewer P cells were injected in higher order infections, and thus bacteria needed more time to replicate and reach sufficiently high numbers to kill the larval host (Fig. S2).

### The most virulent species is also more abundant both in mono and mixed infections

We then asked what the underlying reason could be for our observation that the most virulent species drives host survival dynamics. One explanation would be that the more virulent species grows better in the host, making it more abundant and thereby exerting a stronger effect on the host. We tested this hypothesis in mono infections first and found that bacterial load (CFU per larva) of the four pathogens followed the exact order of their virulence at 12 hours post infection (hpi, see Fig. S3 in the supplemental information, and Table S3 for the full statistical analysis). At the earlier timepoint (6 hpi), the same pattern holds for B, C, and P, whereas K had much higher CFU inside the host than expected from its virulence. The data imply that K initially grows well in larvae, while it is compromised later during the infection. Overall, the mono infection data suggest that pathogen load in a host positively links to its level of virulence (i.e., negatively with host survival).

Next, we explored whether the same is true for mixed infections, i.e., whether the dominant effect of the more virulent species might be caused by its higher abundance in a co-infection. Indeed, in all cases the more virulent species was also the more abundant species at 12 hpi (Fig. 3). For example, P as the most virulent of our pathogens dominated in terms of abundance in all larvae in any of its mixes at 12 hpi. However, we also found evidence for more complex dynamics, where relative species abundance changed over time. Especially in the infection pairs K+P and K+B, we observed that K dominated at 6 hpi despite being the less virulent species, a pattern that disappeared at 12 hpi. Important to note is that in five out of six pairings both pathogen species coexisted in the host during the course of the infection in a large proportion of larvae.

**Figure 3.**
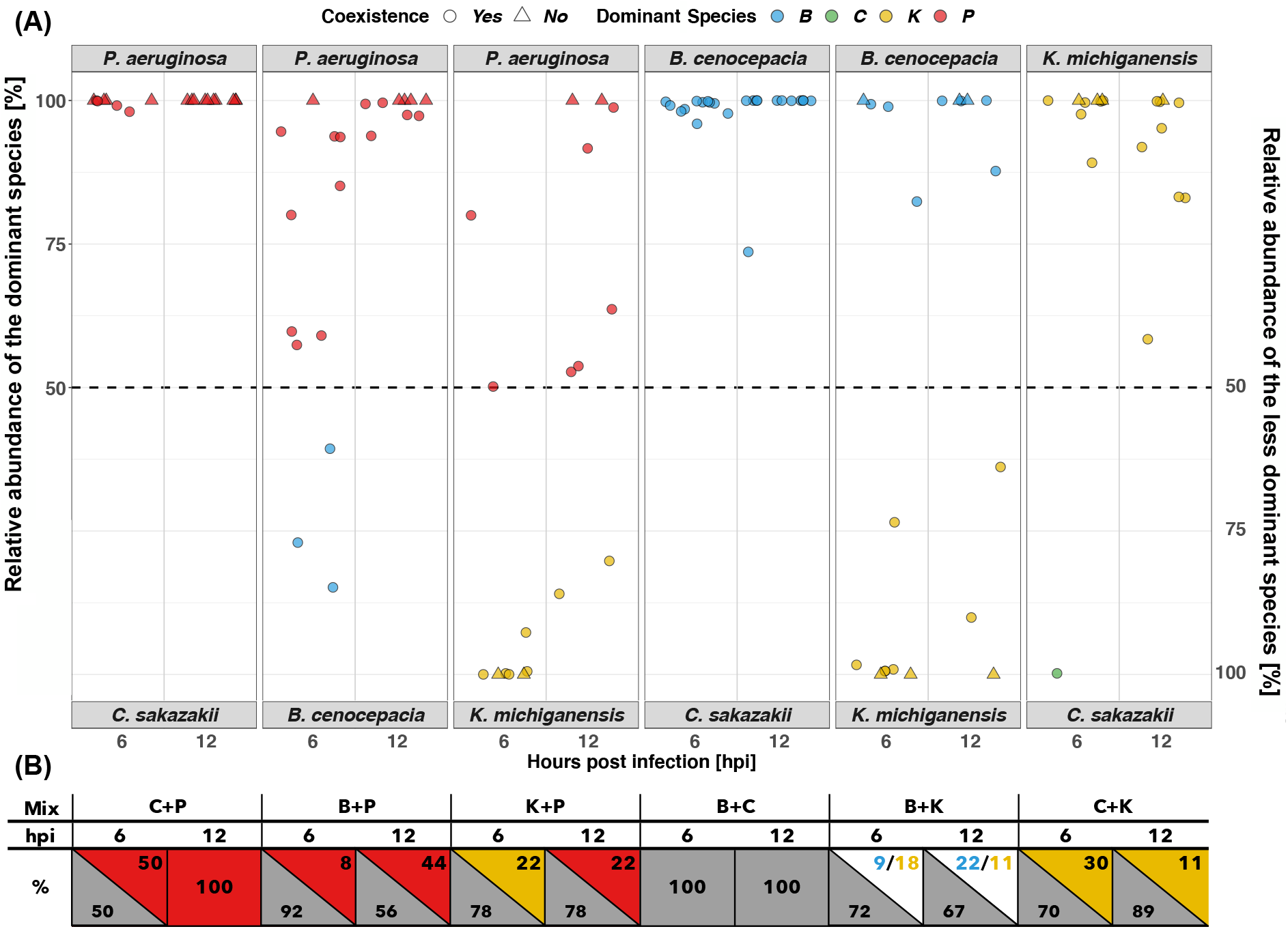
The more virulent pathogen is more abundant in pairwise infections and coexistence can occur in five out of six pairings throughout the infection. (A) Relative abundance in pairwise infections of *G. mellonella* larvae at 6 and 12 hours post infection (hpi). The panels follow the order of the more virulent species in the mix, from left to right, with the more dominant species being labeled on top of each panel. The left and right y-axes show the relative abundance of the more dominant and less dominant species, respectively. Each datapoint represents one larva and is colored according to the more abundant pathogen species found in the larva. Circular datapoints represent pairwise infections in which both pathogens were found at 6 hpi or 12 hpi and triangles indicate datapoints for which only one of the injected species remained in the host (or the rarer species was below detection limit). (B) Rectangles and triangles show the percentage of larvae in which either coexistence (grey) of pathogens occurred or only a single species (color) was left. Data are shown for all pairwise infections both at 6 hpi and 12 hpi. Data are from 4-5 individual experiments with 2-3 larvae per treatment, resulting in a total of 8-12 larvae per treatment.

### Negative interactions dominate in pairwise infections

We used the CFU data from mixed infections to test whether the growth of any of the four species was influenced by the presence of a co-infecting species across two time points (at 6 and 12 hpi). We found that the co-infecting species had either no effect or a negative effect on a focal species in a pathogen-pair specific way (Fig. 4, see Table S4 for the full statistical analysis). The negative effect was either independent of how prevalent the co-infecting species was (main effect) or stronger with increasing CFU of the co-infecting species (CFU effect).

**Figure 4.**
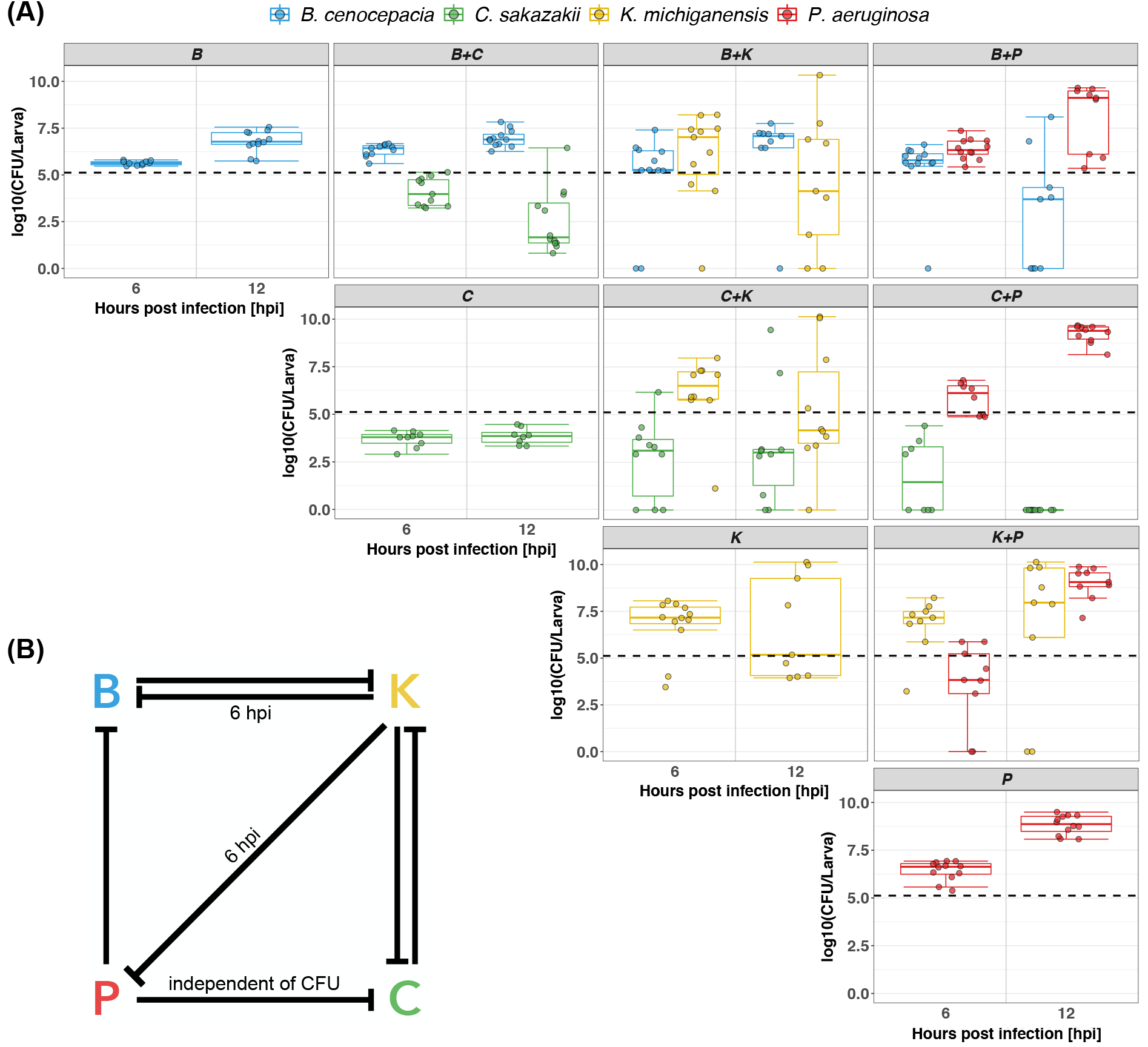
Bacterial load in *G. mellonella* larvae infected with a single pathogen or pairs of pathogens 6 and 12 hours post infection (hpi). (A) Boxplots depict the number of CFU per larva for the mono infections of B, C, K, and P, and all pairwise infections for the two time points measured. Dots represent data from individual larvae. The dashed black line shows the injection dose of 50 000 CFU per species in mixed infections. The dose for mono infections was 100 000 CFU. To be able to directly compare the CFU between mono and mixed infections, we halved the actual CFU counts obtained from the mono infections. Boxplots show the median (line within the box) with the first and third quartiles. The whiskers cover 1.5x of the interquartile range or extend from the lowest to the highest value if all values fall within the 1.5x interquartile range. Data are from 4-5 individual experiments, each featuring 2-3 larvae per treatment, resulting in a total of 8-12 larvae per treatment. (B) Interaction network of our bacterial consortium based on the statistical analysis of the data for both timepoints unless indicated otherwise. Significant negative impact of one species on another is depicted by a stop arrow. The label “independent of CFU” indicates that this effect was not determined by the amount of the co-infecting pathogen.

*B. cenocepacia* (B): The growth of this pathogen increased from 6 hpi to 12 hpi (t_77_ = 2.66, p = 0.0095), and was not affected by C (t_77_ = −0.10, p = 0.9171). In contrast, B growth was compromised by increasing numbers of K at the earlier timepoint and P at both timepoints (CFU effect of second species, for K: t_77_ = −4.73, p < 0.0001; for P: t_77_ = −4.17, p < 0.0001). The presence of P at 12 hpi resulted in a CFU drop of B even below the inoculum size in many larvae.

*C. sakazakii* (C): In mono infections, this pathogen’s CFU dropped below the inoculum size at both timepoints, suggesting that it cannot replicate within the host. While the presence of B did not affect C at any given time, both K and P significantly compromised C abundance (main effect, for K: t_66_ = −3.29, p = 0.0016; for P: t_66_= −2.20, p = 0.0314).

*K. michiganensis* (K): In line with the host survival data (Fig. 1A and B), we found more CFU of this pathogen the older the larval host was (t_71_= 4.27, p < 0.0001). The growth of K significantly decreased with higher CFU of the co-infecting B at both timepoints (CFU effect of B: t_71_ = −2.48, p = 0.0156). Interestingly, our model predicts that K reaches higher CFU if C is present at sufficiently high numbers (CFU effect of C: t_71_ = 2.90, p = 0.0050), but because C never reached high CFU itself, the overall effect of C on K was negative in our experiments (main effect of C: t_71_ = −2.92, p < 0.0047).

*P. aeruginosa* (P): The CFU of P significantly increased from 6 to 12 hpi (t_73_ = 9.89, p < 0.0001). P growth was only affected by K, which had a strong negative impact at 6 hpi (CFU effect of K: t_73_ = −4.40, p < 0.0001).

While the number of interactions decreased over time (58% at 6 hpi; 42% at 12 hpi), all interactions were negative at both timepoints. Our four bacterial species spanned from being inhibited by one (P) or two other species (B, C, K) and inhibiting one (B, C), two (P) or three (K) other species (Fig. 4B). At 12 hpi, the competitive strength of the four pathogens followed their order of virulence and growth, whereby P suppresses B and C; B suppresses K; and K and C mutually repress each other with the former being slightly stronger.

### Targeting the more virulent species in mixed infections increases host survival

The main conclusion of the above results is that there are positive associations between a pathogen’s growth inside the host, its competitiveness towards other pathogens, and its virulence. Consequently, we predicted that treating the most virulent species in a multispecies bacterial infection would be an effective strategy to increase host survival. To test this prediction, we co-infected *G. mellonella* larvae with B+P and compared host survival of the mixed infection to the mono infections of B and P, with all conditions receiving either no treatment or a single dose of gentamicin after two hours. Given that B is inherently resistant to gentamicin, we predicted that the more virulent pathogen P should be selectively killed and the survival dynamics of the mixed infection should follow the one of the B mono infection.

In the absence of gentamicin, we recovered our previous findings (Fig. 2B) that mixed B+P infections followed the virulence trajectory of P mono infections (Fig. 5). In stark contrast and in support of our prediction, we observed that host survival in B+P mixed infection more closely followed the trajectory of the B mono infection under gentamicin treatment (Fig. 5). This finding suggests that P was selectively removed by gentamicin, which is supported by our mono infection data showing that larval survival is very high for the P infection treated with gentamicin. Overall, B+P infections treated with gentamicin were significantly less virulent than B+P infections not receiving gentamicin treatment (log-rank test: B+P+gentamicin vs. B+P: p < 0.0001). These results support the view that treating the most virulent species in a polymicrobial infection is a promising strategy to reduce host mortality.

**Figure 5.**
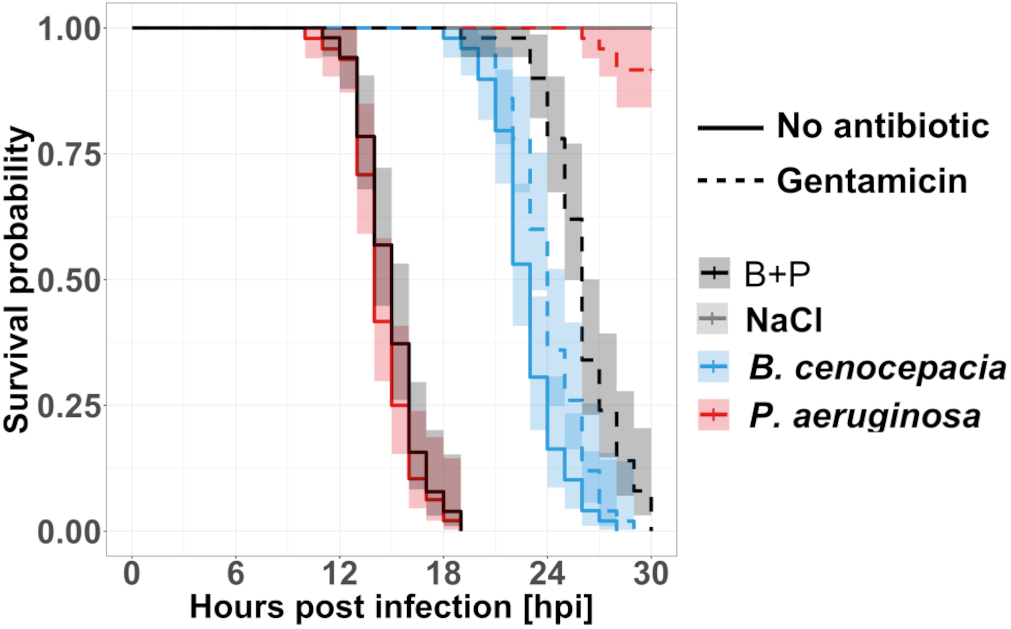
Treating the more virulent pathogen reduces host mortality in mixed infections to the level of the less virulent pathogen. Kaplan Meier survival curves of *G. mellonella* larvae injected with either NaCl (control, grey), B (less virulent pathogen, blue), P (more virulent pathogen, red), or B+P mixes (black). Larvae received either a gentamicin treatment (dashed lines) two hours after the bacterial/NaCl injection or no antibiotic treatment (solid lines). Survival curves are based on a total of 48-54 larvae per treatment from three independent experiments.

## Discussion

There is increasing evidence that polymicrobial infections are common (1–4) and that ecological and evolutionary interactions between co-infecting pathogens can affect host morbidity (5–7). In our study, we examined whether there are generalizable patterns that characterize interaction dynamics between pathogens and a common host in multispecies infections. For our experiments, we used the larvae of *G. mellonella* as the host and infected it with four opportunistic human bacterial pathogens alone and in all possible combinations of mixed infections. We tested the rank order of virulence for the four pathogens and found that it was identical to the rank order of growth in the host. In addition, the more virulent species in mono infections were also better at outcompeting other species in mixed infections. A consequence of this was that the most virulent species determined host survival dynamics in mixed infections regardless of the number and type of pathogens mixed. Our findings, which held for all pathogen combinations tested, reveal an infection dynamic that is not covered by any of the current models for pathogen virulence in mixed infections.

While we found that the virulence of a specific pathogen is positively linked to its growth and competitiveness in the host, our co-occurrence analysis could give us an idea of the relative importance of pathogen growth versus competitiveness and how it varies across pathogen species combinations (Fig. 4). In the pathogen interaction network (Fig. 4B), the absence of an interaction suggests that differences in growth dominate a co-infection, meaning that the faster growing species simply outperformed the slower growing one. This was the case for five (6 hpi) and seven (12 hpi) out of the total twelve interactions. Conversely, the remaining seven (6 hpi) and five (12 hpi) interactions that were negative would imply that competitiveness played a more prominent role. At least two types of competition could be involved. First, the faster-growing species limits resource availability for the slow-growing species, reducing its growth and survival due to starvation. Second, the more competitive species secretes a toxin or deploys contact-dependent mechanisms to target and kill the less competitive species in interference competition. While our results do not allow to differentiate between resource and interference competition (34), it is likely that both mechanisms matter. For example, in our mixed infections with K+B, and K+P, we observed that K inhibits its competitors early on during the infections (6 hpi) but is outcompeted at a later stage (12 hpi). One option is that the inhibitory effects early during the infection could be explained by K secreting a toxin, while the dominance of B and P later in the infection could be due to resource competition advantages. Alternatively, it could be that K grows quickly but inefficiently while its competitors (B and P) grow more slowly but efficiently and thus outpace K over time.

Important to note is also that all pathogen interactions were either negative or neutral, but never positive. This finding supports the view that competition is much more prevalent between bacterial species than positive interactions, where one species unilaterally or mutually benefits another species (35, 36). The prevalence of competitive interactions is perhaps expected given that the pathogens interacted in a closed environment (i.e., the larva), where both resource availability and host longevity are limited (37).

Competition between pathogens is also a major component of mathematical models predicting virulence levels (38). A traditional set of models assumes that genetically diverse pathogens engage in increased levels of resource competition in mixed infections, which is predicted to exacerbate virulence (37). Other models examined the effect of infighting between pathogens through toxin secretion (39) or competition for publicly shared virulence factors (40). These models predict that increased competition between pathogens should decrease virulence in mixed infections. Empirical support for these models vary, lending support to both types of predictions (41–44). A key insight from these studies is that the biological details of pathogen interactions matter. While our study was not designed to test specific model predictions, our results put forth a third virulence scenario, namely that pathogen interactions in mixed infections neither increase nor decrease virulence, but the virulence trajectory simply follows that of the more harmful species. A first example of this scenario was reported by Massey et al. (43) and here we reveal its generality. As discussed above, this pattern can arise when pathogen traits that are relevant for infections (virulence, growth, and competitiveness) are positively connected with one another. This virulence model could be particularly relevant for infections with opportunistic pathogens that co-incidentally end up in the same host with no prior interaction and coevolution history (3).

Our data further indicate that host factors also influence pathogen virulence patterns. For example, we found that younger larvae had longer survival times than older larvae when infected with B, C, and K. Given that *G. mellonella* has an innate immune system similar to the one of vertebrates, our results indicate that the immune response works well against weaker pathogens (like C and K) but deteriorates with age (i.e. number of days after the arrival of the larvae in our laboratory). Another host effect we observed is that *G. mellonella* managed to control infections of C and K at low injection doses, again highlighting the potency of the insect’s immune system. A further indication that pathogen performance is modulated by the host stems from an additional experiment, which we conducted with a medium that mimics the hemolymph of the insect order Lepidoptera (Fig. S4). Although this medium nutritionally matches insect conditions, we found the rank order of pathogen species growth to be markedly different compared to growth in live hosts (insect condition *in vitro*: C > P > K > B vs. *in vivo*: P > B > K > C). Clearly, host effects are important and likely feedback on pathogen-pathogen interactions, which is why they should be considered in future work on polymicrobial infections.

In conclusion, we can draw two generalities from our four-bacteria-infection system. First, no matter what pathogen combination we used, the most virulent pathogen dictated host survival. This led us to predict that targeting the most virulent pathogen seems a promising strategy to reduce host morbidity/mortality in polymicrobial infections. We found support for this prediction in a proof-of-principle experiment, where we selectively targeted the more virulent species in a mixed infection with antibiotics. Second, more virulent pathogens grow better in the host and are better in competition with less virulent pathogens. These observations suggest that the same traits (or co-regulated traits) responsible for attacking the host (e.g., virulence factors) could promote pathogen growth and help in competition with other pathogens. Identifying these traits could give rise to promising strategies to control polymicrobial infections.

## Materials and Methods

### Bacterial strains & host

We used the following four opportunistic human pathogens: *Pseudomonas aeruginosa* PAO1 (45), *Burkholderia cenocepacia* K56-2 (46), *Klebsiella michiganensis*, and *Cronobacter sakazakii* (ATCC29004). We purchased the larvae of the greater wax moth *G. mellonella* in their last instar stage from a local vendor (Bait Express GmbH, Basel, Switzerland). We contacted the vendor who confirmed that their larvae had not been treated with any antibiotics. We can therefore rule out the possibility that pathogen interactions in our experiments are affected by residual antibiotic concentrations in the larvae. Upon arrival, we stored them at 4-8° C without food. We considered larval age as the number of days after arrival in our laboratory, since the exact age of larvae was unknown to us. Hence, we only investigated larval age as a relative measure. When examining larval age, we compared larvae that were ordered at the same time (i.e., same batch), but were infected at different time points. To put host age into a wider context, the *G. mellonella* life cycle comprises four developmental phases: egg (3-30 d), larva/caterpillar (6-7 weeks), pupa (6-55 d), and adult (47). We only worked with the second phase, i.e., the larva, at its last out of seven larval stages before turning into a pupa (last instar stage) (48).

### Culturing conditions of bacteria

Bacterial stocks were kept in 25% glycerol and stored at −80° C. For all experiments, we grew bacteria overnight until stationary phase in 5 mL lysogeny broth (LB) in 50 mL Falcon tubes at 37° C and at 170 rpm with aeration (Infors HT, Multitron Standard Shaker). For all species, we centrifuged 1 mL overnight-cell culture in a 1.5 mL Eppendorf tube at 7’500 rcf for 5 min (Eppendorf, tabletop centrifuge MiniSpin plus with rotor F-45-12-11) and washed cultures three times with a 0.8% NaCl solution to remove all original media. Next, we measured the optical density at 600 nm (OD_600_) of the washed cell culture (Amersham Biosciences, Ultrospec 2100 pro spectrophotometer), adjusted it to OD_600_=1, and diluted each species individually to obtain similar cell numbers per milliliter for all species.

While all our main experiments were carried out in the *G. mellonella* larvae, we conducted one additional *in vitro* experiment. Specifically, we grew all four pathogens as monocultures in a medium that mimics the hemolymph of Lepidoptera – the insect order to which *G. mellonella* belongs – called Grace’s insect medium (GIM, Fig. S4). The aim of this experiment was to show that our model host directly affects pathogen growth and is not just serving as a sack to conduct growth assays. Thus, we expected pathogen growth to be different inside the host compared to the *in vitro* GIM environment. We prepared bacterial overnight cultures as stated above to obtain washed and OD_600_=1 adjusted cell cultures. A 96-well cell culture plate (Eppendorf, non-treated, flat bottom) was then filled with 190 μL GIM (Gibco, GIM 1X, supplemented) and 10 μL cells to reach a starting OD_600_=0.001 per well. To reduce evaporation the inter-well spaces and the outer moat space of the microplate were filled with sterile water (13 mL in total). We incubated the microplate at 37° C in a microplate reader (Tecan, Infinite MPlex, monochromator optics) and measured OD_600_ every 15 min for 24 h, shaking for 60 sec before every read. The plate layout was randomized between experiments. All chemicals were purchased from Sigma Aldrich, Switzerland, unless indicated otherwise.

### Infection experiments

Prior to infections, we sorted larvae and evenly distributed them across treatments according to size. Their weight ranged from 278-731 mg with an average weight of 499 mg. We put larvae on ice in a petri dish to immobilize them. Next, we surface-sterilized larvae with 70% ethanol and injected them between the posterior prolegs. We used a sterile hypodermic needle (Braun, 0.45 x 12 mm BI/LB, 26G), a sterile syringe (Braun, 0.01-1 mL Injekt-F Luer Solo), and a programmable syringe pump (New Era Pump Systems Inc, model NE-300) to standardize injection speed. With a flow rate of 2 ml/h, we either injected 10 μL control or bacterial solution (both in 0.8 % NaCl) into larvae within 18 s. We had a second control treatment, where larvae were not injected. The injection dose ranged from 10^2^ to 10^6^ CFU for mono infections and was kept at 10^5^ CFU for experiments that included multispecies infections. In multispecies infections, we mixed equal amounts of each species. The order of treatments was randomized between experiments. For all experiments, we confirmed the injection dose in triplicate by CFU on 1.5% LB agar plates. We distributed injected larvae to individual wells of a 12-well plate for incubation at 37° C in the dark without food. We monitored survival of every larva in regular intervals (hourly during 12-24 hours post infection (hpi) and every two hours during 36-48 hpi). Larvae were considered dead when they did not move upon touch with a pipette tip.

We used gentamicin to selectively target P in co-infections with B. First, we determined the gentamicin concentration that killed P but had no effect on B *in vitro*, since B is inherently resistant to gentamicin (up to a certain concentration). We found that 8 μg/mL gentamicin fulfilled the condition. Thus, we injected 10 μL of a 400 μg/mL gentamicin solution into the larvae, which should result in an internal gentamicin concentration of 8 μg/mL based on the average larval weight (assuming 499 mg ~ 500 mL according to (49)). The gentamicin treatment was applied two hours after the bacteria had been injected so that they had time to establish an infection in the host. To control for any impact caused by the second injection, half of the NaCl sterile control larvae also received the gentamicin treatment.

### Bacterial load in the hemolymph

We determined the bacterial load in all larvae that received mono-, mixed-, NaCl, and no infection at two time points, at 6 and 12 hpi. Since larvae had to be sacrificed during this process, we had two independent batches of larvae that were used for the respective timepoints. First, we placed larvae individually in 2 mL Eppendorf tubes and submerged those in ice until larval movement halted. Then, we opened each larva with a surgical blade (Aesculap AG, Germany) behind the posterior prolegs, with the cut length spanning half of the body width. This procedure is similar to the one used by Harding et al. (50), McCloskey et al. (51), and Admella & Torrents (52). We sterilized the blade with 70% ethanol between different larvae. Next, we drained 10-20 μL hemolymph into a fresh 1.5 mL Eppendorf tube by gently squeezing the larva. The collected hemolymph was mixed and stored on ice. To enumerate bacterial CFU, we serially diluted 10 μL of the collected hemolymph in 0.8% NaCl up to 10^−6^ and plated the appropriate dilutions on LB-agar plates (1.5% agar) in duplicates. The appropriate dilutions varied between 10^0^ (undiluted) and 10^−6^ and depended on the bacterial species. We incubated the plates at 37° C overnight and then manually counted CFU. Because *B. cenocepacia* grew slower than the other species, plates were incubated for two days. Plates with *C. sakazakii* were kept at room temperature for an additional day following overnight incubation for colonies to develop their characteristic yellow color. All four species could be distinguished from each other based on their different colony morphologies (see Fig. S5). We considered plates with a minimum of 25 CFU up to a number of CFU for which we could still distinguish single colonies with confidence. Plates with less than 25 CFU were only considered if none of the plated dilutions from the same individual adhered to this threshold. For mixes where one species occurred at very low frequency (mostly mixes with P), we plated the undiluted hemolymph to see whether the rare species was present at all. While both C and K were visible even in a lawn of P, this was not true for B. For this species combination, we cannot conclude that B was completely absent in certain larvae but rather that P was highly dominant. We calculated the CFU/larva by multiplying the volume of plated hemolymph with the average larval weight since larval weight is almost identical to larval liquid volume according to Andrea et al. (49).

In approximately 20% of the extracted larval hemolymph, we detected bacteria different from our four focal species. We determined the identity of ten representative isolates by 16S rRNA sequencing (colony PCR using standard primers 1492r and 27f). We identified *Enterococcus casseliflavus*, and *E. gallinarum*, two common insect gut bacteria (53, 54), as the closest relatives of our isolates with sequence identities of 89-99%. These results suggest that we had drained certain larvae too extensively, so that we not only collected bacteria from the hemolymph but also from the gut. For this reason, we excluded these larvae from all further analysis.

### Statistical analysis

All statistical analyses were performed with R (version 4.1.1) and the interface RStudio (version 2021.09.0) (55). For the dose-response curves in Fig. 1A, we measured mean survival time of the host as the area under the full survival curve using the function survmean from the survival package (56). To build a log-logistic model we used the drm function from the drc package (57). To compare host survival in mono versus mixed infections, we built a separate model for each panel shown in Fig. 2, comparing host survival of a particular pathogen combination. The statistical results were robust across three different methods comparing host survival: the non-parametric log-rank test, the semi-parametric Cox proportional hazards model, and the parametric Weibull regression. See Table S2 for a model comparison. Non-parametric log-rank tests were also used to compare host survival between mono and mixed infections with or without gentamicin treatment (Fig. 5, Table S2).

To determine whether the pathogens significantly vary in their bacterial load in mono infections, we compared the CFU extracted from the hemolymph of host individuals at 6 hpi and 12 hpi. We chose a linear model using generalized least squares to account for differences in variance between the species, which was mainly caused by *K. michiganensis* at 12 hpi (see Table S3 for the full statistical analysis). Next, we performed a post-hoc analysis using the Tukey honest significant difference test with p-value adjustment to compare the bacterial load between species.

To compare bacterial load in the hemolymph in mono-versus mixed infections, we built a separate linear model for each pathogen to test whether its growth was influenced by the presence of another species, the CFU of this other species and time (at 6 hpi versus 12 hpi). Since the injection dose was always the same independent of how many species were injected, we compared for each species half the CFU found in mono infections to the total CFU (for the focal species) observed in pairwise infections. To obtain normally distributed residuals, we transformed all CFU values in our models. We used the function transformTukey from the rcompanion package to find the best transformation (58). Non-significant interaction terms and main effects were removed from models until a minimal model was obtained. Most of the minimal models included both a main effect of the co-infecting species and its CFU effect. We often found that these two explanatory variables had opposing signs, which means that the influence of the co-infecting species on the focal species depended on the actual CFU of the co-infecting species. For this reason, we assessed the threshold CFU of the co-infecting pathogen at which its effect on the focal pathogen switches from being positive to being negative. This was done for each pathogen combination. We could then compare this CFU threshold to our experimental CFU values to check whether a co-infecting species stimulated or inhibited a focal species. To calculate the standard error for confidence intervals of this threshold and the p-value (one sample t-test versus zero) we used the Taylor expansion in the R package propagate (59).

## Data availability

All raw data sets will be deposited in the figshare repository (LINK).

## Supplementary information

Supplementary information will be made available online: Supplemental file, XLSX file, XX MB.

## Acknowledgements

We thank Kayla King, Anna-Liisa Laine, Alex Hall, and Roland Regoes for their scientific inputs. We also thank Nadine Koch for showing us how to conduct injections with the larvae of *G. mellonella*.

## Compliance with ethical standards

Conflict of interest: The authors declare that they have no conflict of interest.

