## Supplementary figures for "Negative interactions and virulence differences drive the dynamics in multispecies bacterial infections"

This file contains supplementary information for the article

The supplementary information comprises:

- 5 supplementary Figures: Figure S1 - S5

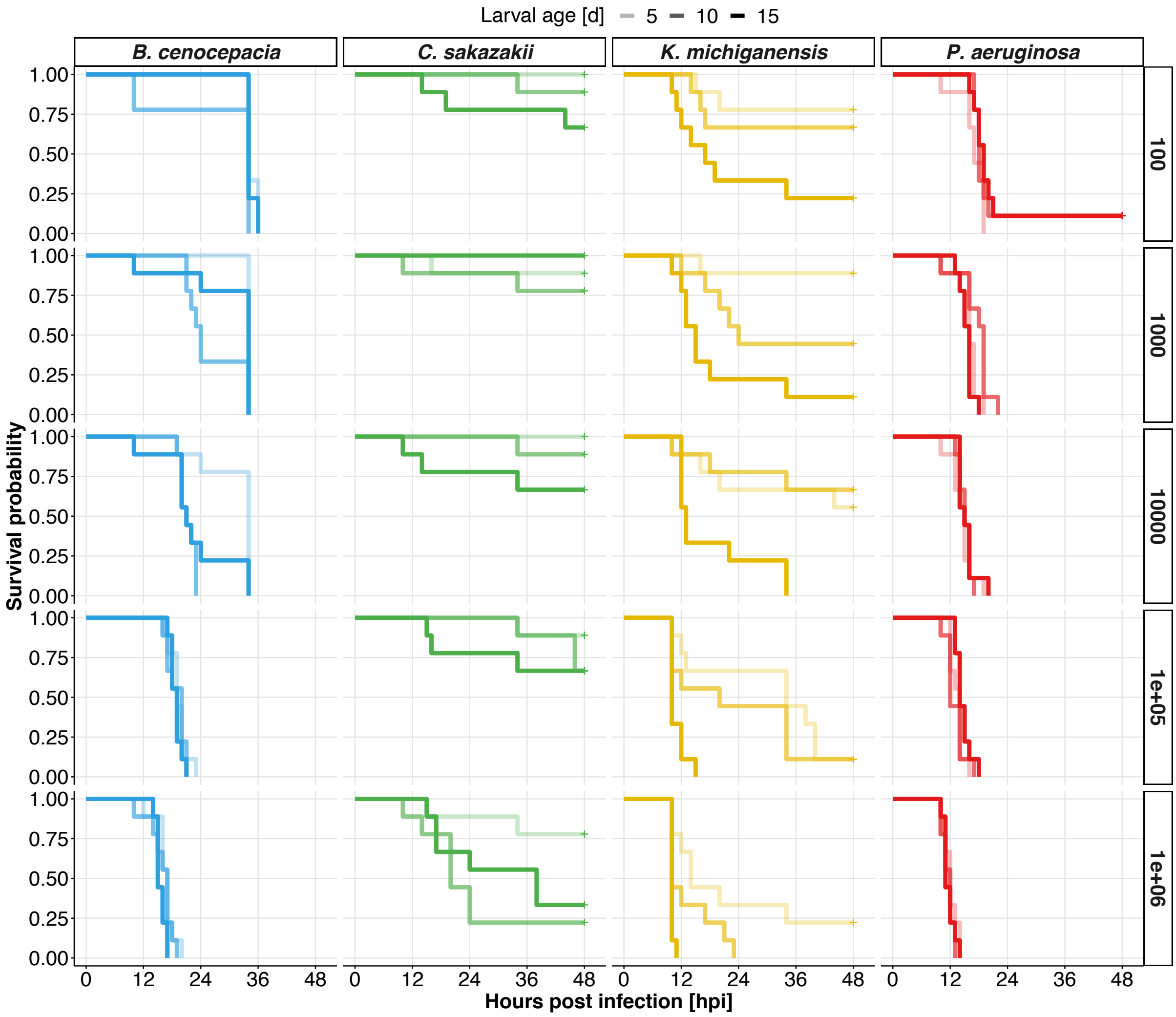


**Figure S1.** Virulence significantly differs between our four bacterial pathogens and increases both with higher injection doses for all species and host age for K, C, and marginally for B but not for P (log-rank: $p < 0.05$). Kaplan Meier survival curves are shown for each of the four bacterial pathogens. Larvae were injected with pathogens across a range of doses, from 100 to 1 million CFU (secondary y-axis on the right). Relative larval age ranged from 5 to 15 days old (i.e., days after the arrival in our laboratory), indicated by increased line opaqueness. Data are from three individual experiments with a total of 30-36 larvae per treatment.

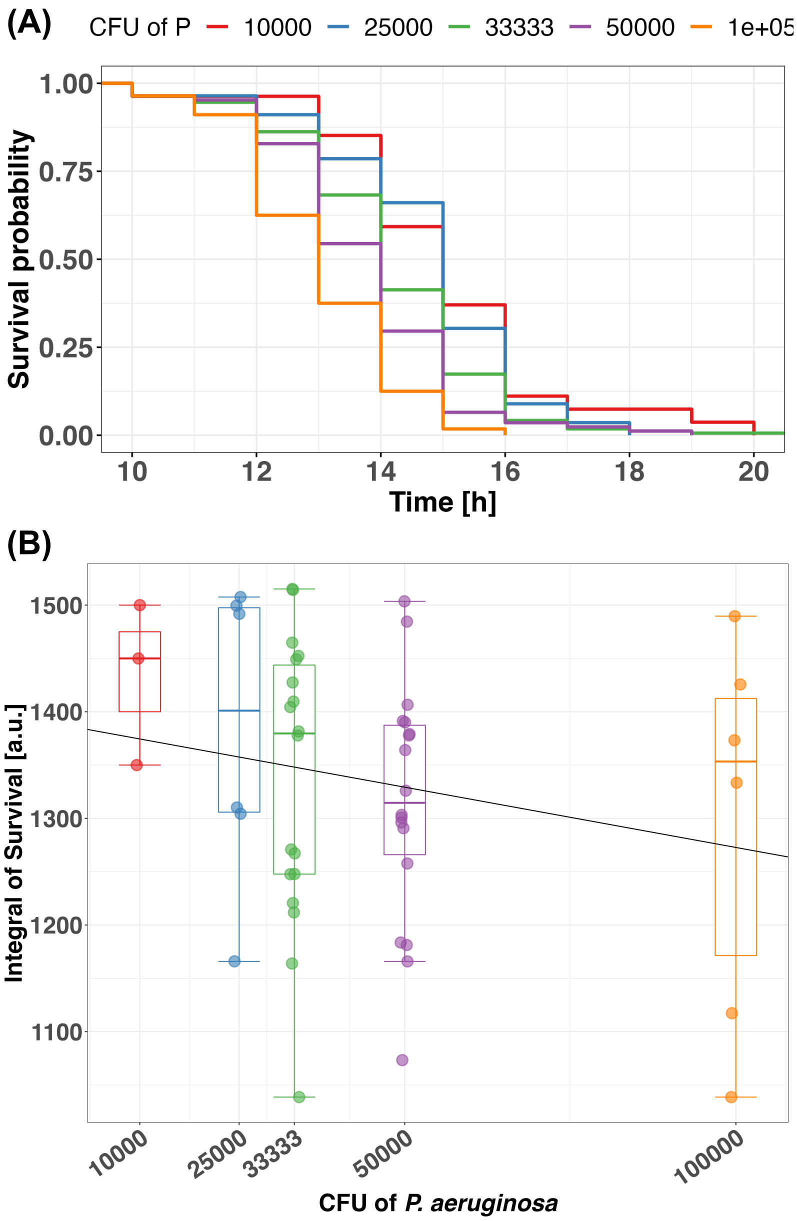


**Figure S2.** A higher infection dose of *P. aeruginosa* (P, expressed as CFU) leads to decreased survival of *G. mellonella* larvae across mono and mixed infections. Our main findings suggest that host survival is determined by P, whenever this most virulent species is present in the co-infection regardless of the identity of the co-infecting pathogen. However, because the P infection dose varies between pairwise (50,000 CFU), triple (33,333 CFU), and quadruple (25,000 CFU) infections, we predict variation in host survival to lay between the P mono-infections with 10,000 CFU (highest host survival) and CFU 100,000 (lowest host survival). Our analysis supports this prediction. **(A)** Kaplan Meier survival curves show a stepwise decrease of host survival in response to higher P infection doses. Pairwise infections comprise B+P, C+P, and K+P mixes; triple infections comprise B+C+P, B+K+P, C+K+P mixes; the quadruple infection comprises the B+C+K+P mix. Data are from 3-5 independent experiments with 10-12 larvae per treatment. Due to the unequal number of combinations, the mono and quadruple survival curves are based on 30-60 larvae, while the pairwise and triple infections are based on 150-180 larvae. **(B)** Negative correlation between P infection dose and larval survival. We derived the integral of survival from the area under the full survival curve for each independent experiment (shown as dots) separately using spline fits. The data show the expected negative correlation but also considerable variation across experiments (although the range of values all represent steep killing curves characteristic of P). Boxplots depict the median (line within the box) with the first and third quartiles. The whiskers cover 1.5x of the interquartile range or extend from the lowest to the highest value if all values fall within the 1.5x interquartile range. The regression line was calculated based on a linear model with the integral of survival as response variable and the *P. aeruginosa* infection dose (CFU) as a continuous explanatory variable.

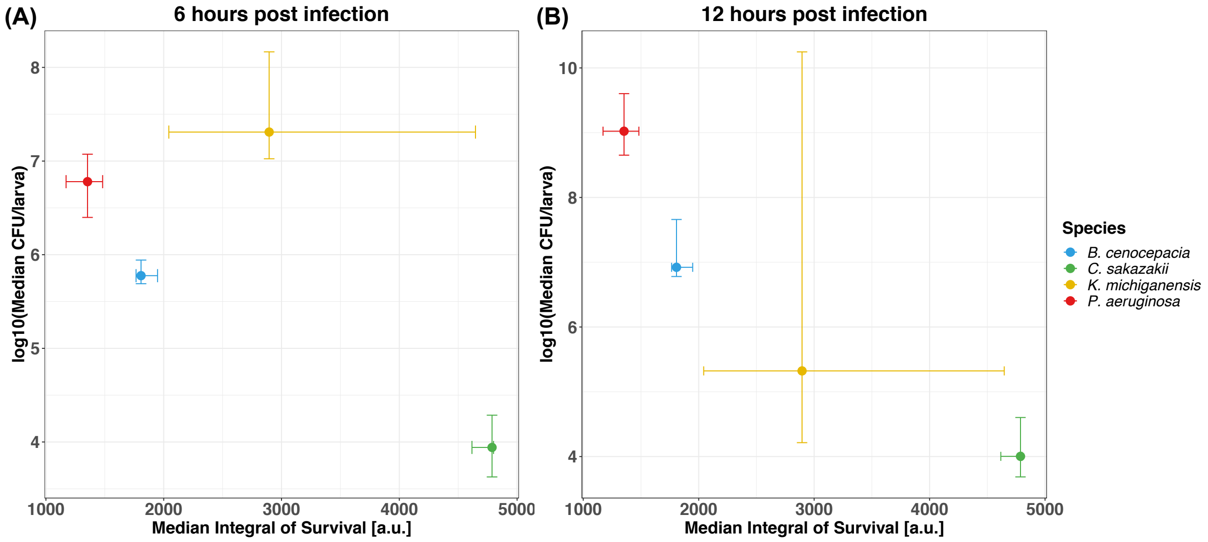


**Figure S3.** Bacterial load within the host positively links to virulence at later stages of the infection. Associations between median survival and median pathogen load are shown for (A) at 6 hours post infection (hpi) and (B) at 12 hpi. The datapoints represent the median values and the whiskers cover the range between the first and third quartiles. The median integral of survival was measured as the area under the full survival curve, and stems from five experiments, each with 10-12 larvae per treatment, resulting in a total of 50-60 larvae per treatment. The median CFU/larva values originate from 4-5 individual experiments with 2-3 larvae per treatment each, resulting in a total of 8-12 larvae per treatment.

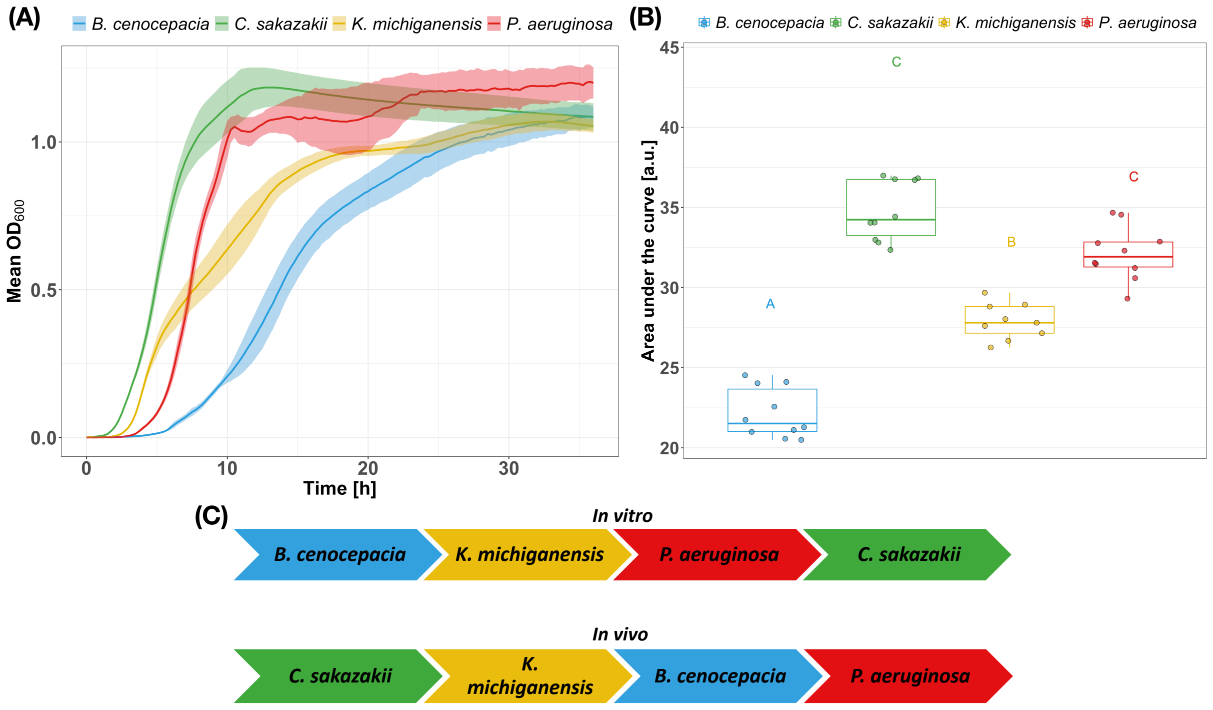


**Figure S4.** Rank order of growth of the four pathogens in Grace's insect medium differs from the rank order of growth observed in the larvae of *G. mellonella* (Fig. 4), suggesting that the host environment impacts pathogen performance. We examined bacterial growth *in vitro* in Grace's insect medium (GIM), a medium that mimics the haemolymph of Lepidoptera – the insect order to which *G. mellonella* belongs. (A) Growth curves show the mean $OD_{600}$ of the four pathogens over the course of 24 h in shaken, liquid GIM cultures. Each curve is based on 9-10 replicates from a total of 3 independent experiments (each featuring 3-4 replicates). The dashed, vertical line at 12 h marks the time point at which CFU where enumerated in the host (i.e., at 12 hours post infection, hpi). (B) Statistical comparison of the growth performance of the four pathogens based on the area under the growth curve until 12 h using the gompertz curve fit. Boxplots show the median (line within the box) with the first and third quartiles. The whiskers cover 1.5x of the interquartile range or extend from the lowest to the highest value if all values fall within the 1.5x interquartile range. Different letters above boxplots indicate significant growth differences (alpha = 0.05) between pathogen species based on a linear mixed effect model (p-value for pairwise comparisons adjusted with the FDR method). (C) Qualitative arrow chart comparing the growth rank order in GIM (12 h) to the growth rank order in *G. mellonella* larvae (12 hpi), ordered from the lowest (left) to the highest (right) performing species. The rank orders differ fundamentally. While *C. sakazakii* did not seem to replicate in the larvae, *C. sakazakii* was the fastest growing species in Grace's insect medium, even outperforming *P. aeruginosa*. Conversely, *B. cenocepacia* performed well in the host but was the poorest performer in the insect medium. This comparison suggests that host factors (other than nutrients) had a strong impact on pathogen growth in vivo.

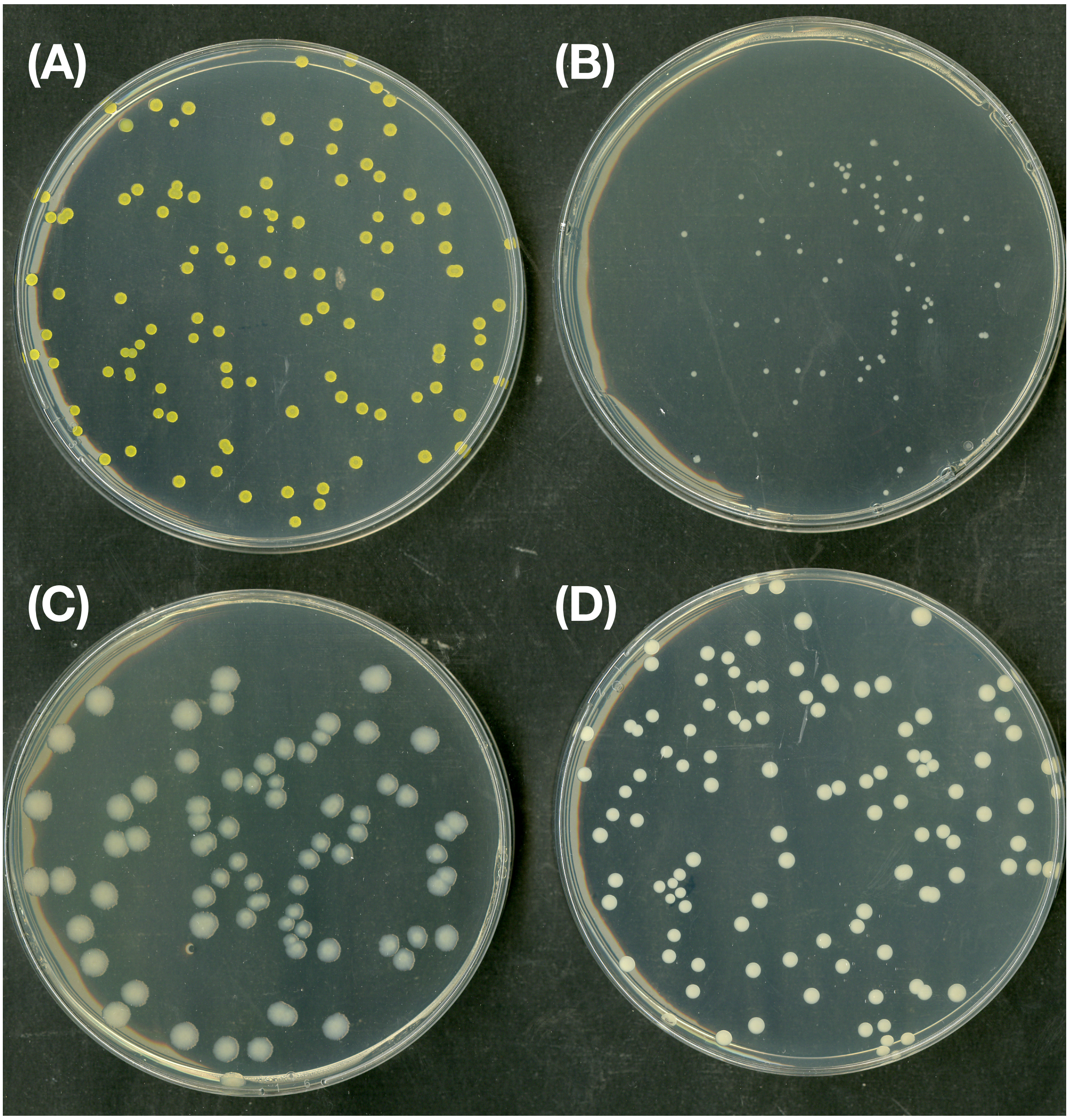


**Figure S5.** The colony morphology of the four bacterial species differs and allows an unambiguous distinction on LB-agar plates. **(A)** *C. sakazakii* has bright yellow colonies **(B)** *B. cenocepacia* has small white colonies **(C)** *P. aeruginosa* has whitish colonies that get more transparent towards the frayed edges **(D)** *K. michiganensis* has beige, perfectly round colonies. Pictures show representative plates.
